# CellWalker: A user-friendly and modular computational pipeline for morphological analysis of microscopy images

**DOI:** 10.1101/2023.02.13.526957

**Authors:** Harshavardhan Khare, Nathaly Dongo Mendoza, Chiara Zurzolo

**Affiliations:** Institut Pasteur, Membrane Traffic and Pathogenesis, Université Paris Cité, CNRS UMR 3691, F-75015 Paris, France; Centro de Investigacion en Bioingenieria - BIO, Universidad de Ingenieria y Tecnologia - UTEC, Lima 15063, Peru

## Abstract

The implementation of computational tools for analysis of microscopy images has been one of the most important technological innovations in biology, providing researchers unmatched capabilities to comprehend cell shape and connectivity. Most available tools frequently focus either on segmentation or morphological analysis, thus not providing an inclusive pipeline. We introduce CellWalker, a computational pipeline that streamlines and connects the segmentation step with the morphological analysis in a modular manner. This python-based pipeline starts with ‘visible-source’ IPython notebooks for segmentation of 2D/3D microscopy images using deep learning and visualization of the segmented images. The next module of CellWalker runs inside Blender, an open-source computer graphics software. This addon provides several morphometric analysis tools that can be used to calculate distances, volume, surface areas and to determine cross-sectional properties. It also includes tools to build skeletons, calculate distributions of sub-cellular organelles. Overall, CellWalker provides practical tools for segmentation and morphological analysis of microscopy images in the form of an open-source and modular pipeline which allows a complete access to fine-tuning of algorithms through visible source code while still retaining a result-oriented interface.

**Availability and implementation:** CellWalker source code is available on GitHub (https://github.com/utraf-pasteur-institute/CellWalker-notebooks and https://github.com/utraf-pasteur-institute/CellWalker-blender) under a GPL-3 license.

## Introduction

One of the most significant technological advancements in biology is the use of computer tools for the processing of microscopy images, providing unprecedented opportunities to understand cell morphology and connectivity, and investigate structure-function relationship. For example, the reconstruction of microscopy images of tissues has facilitated quantification of neuronal structures such as synapses and dendrites, subcellular organization of organelles like mitochondria, endoplasmic reticulum and microtubules (Feng *et al*., 2022; Jorstad *et al*., 2018; Kubota *et al*., 2018; Polo *et al*., 2020; Tamada *et al*., 2017). There are multiple computational tools available that allow the analysis of 3D microscopy data such as Microscopy Image Browser, DeepMIB, VAST, Ilastik (Belevich *et al*., 2016; Belevich and Jokitalo, 2021; Berg *et al*., 2019; Berger *et al*., 2018). Typically, these programs include modules for data preprocessing, automated segmentation, annotation and morphological analysis.

Throughout the past few years, image analysis programs have benefited the study of life sciences. But current technological advances in biology are facing new challenges in terms of accessibility of software tools and the underlying algorithms (Myers, 2012). Most of the commercial software tools are not open-source, which may lead to uncertainty about the working principles and inability to modify the algorithms. On the other hand, many open-source programs come from a unique research question and not readily suitable for different type of investigations, creating a vast number of unused tools (Cardona and Tomancak, 2012), with some notable exceptions such as Biodepot-workflow-builder and ZeroCostDL4Mic (Hung *et al*., 2022; von Chamier *et al*., 2021). Therefore, there is a necessity for open-source programs that include a complete pipeline for processing large microscopy biological datasets and that can be employed across different multidisciplinary biological studies by providing a flexible and modular structure.

Through CellWalker, we aim to provide a set of tools that include both segmentation and morphological analysis in the form of a pipeline that contains independent modules dedicated for various tasks. We utilize existing open-source technologies to make CellWalker visible-source wherever possible. This approach allows users to use customized input data, tweak the algorithms and employ the in-built tools in the software packages inside which the CellWalker modules run.

## Materials and methods

CellWalker is a computational pipeline written in python (version ≥ 3.10) for segmentation and morphological characterization of 2D/3D bioimages (**Fig. 1**). While it is currently developed for cell-biological applications, it may be extended to tissue or organ-level analysis. The pipeline can be divided into two stages-Image Segmentation and Morphological Analysis. There are several steps involved in each of these stages (Please see **supplementary information** and the **GitHub repositories** for more details).

**Figure 1.**
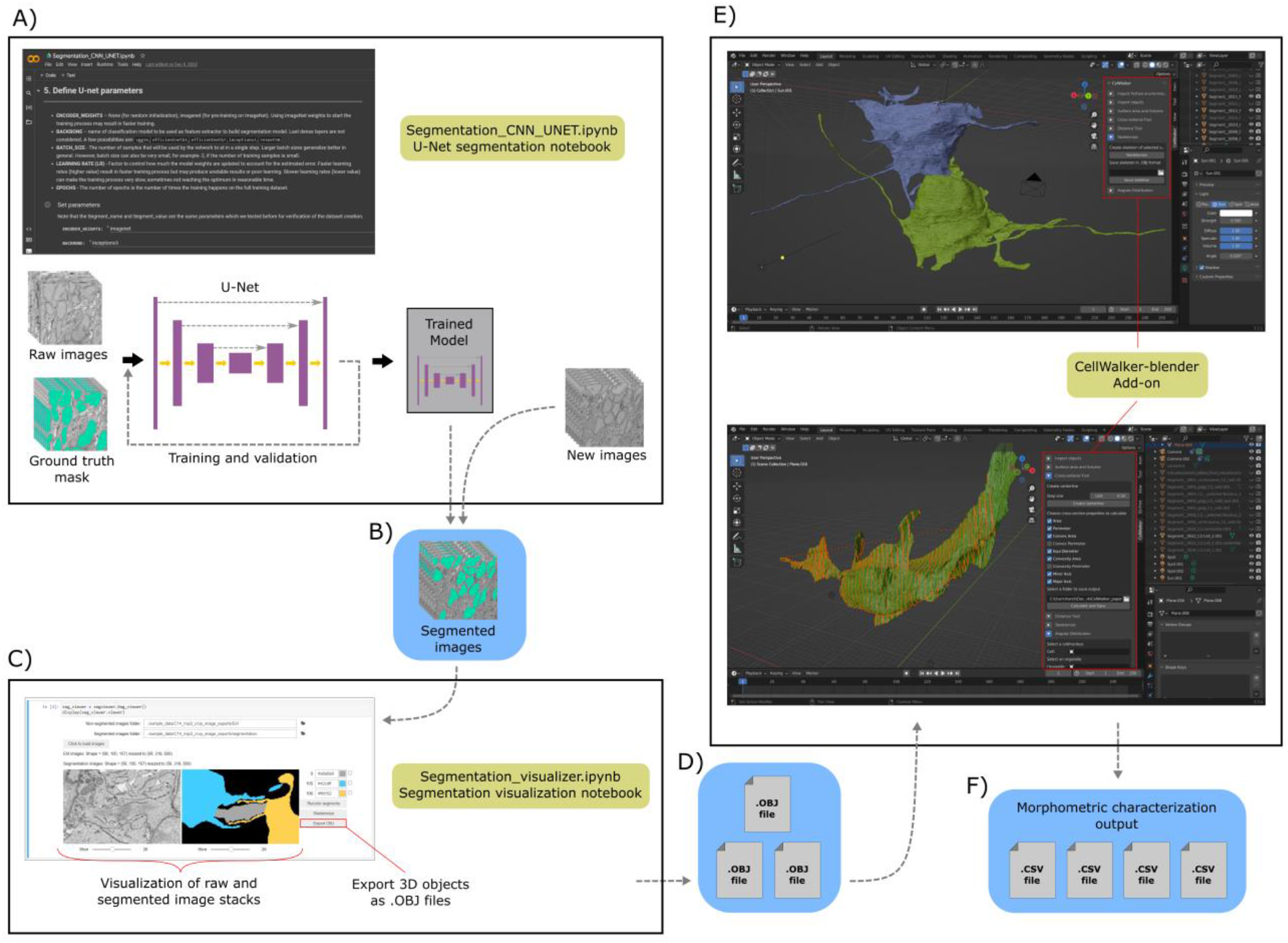
CellWalker pipeline: A) Jupyter-notebook for segmentation using UNET; B) Segmented image stack; C) Jupyter-notebook for visualization of segmentation and exporting 3D objects as. OBJ files; D) Exported. OBJ files; E) CellWalker-blender Add-on inside Blender interface; F) Output of morphometric characterization as. CSV files.

### Image Segmentation

The CellWalker pipeline starts with a module for image segmentation provided as a set of IPython notebooks (**SI Section 1**) that are open-source as well as visible-source, enhancing the flexibility and accessibility for computational biologists while keeping the code execution sufficiently simple for non-coding users. The IPython notebooks can also be freely executed on cloud computing platforms like Google Colaboratory, thus alleviating the need for installation on user’s computer while accessing the data directly from user’s Google Drive.

There are three primary functions of these segmentation notebooks: 1) Automatic segmentation using UNET convolutional networks, 2) Visualization of segmented images and 3) Reconstruction of 3D objects using segmented images.

The automated segmentation notebook (**SI Section 1.1**) implements a 2D UNET algorithm (Ronneberger *et al*., 2015). After training on manually segmented ground truth images, this 2D- UNET is applied to every slice of a 3D image to obtain a 3D segmentation (**Fig. 1A, SI Section 1.1**). The segmentation output is generated as a stack of 2D segmented images (**Fig. 1B**) which can then be supplied to the segmentation viewer notebook for visualization. The segmentation viewer (**Fig. 1C, SI Section 1.2**) can display original and segmented images side-by-side (Haase *et al*., 2021) for direct confirmation of the quality of segmentation. It is possible to set colors of the segmented objects to assist in visualization of multi-object segmentations. The segmentation viewer also provides a function to transform voxelated 3D segments into 3D meshes using a marching cubes algorithm. These meshes are exported as 3D models in a generic Wavefront OBJ format (**Fig. 1D, SI Section 1.2**) which is compatible with other modeling and simulation programs including Blender and 3Dsmax.

Biologists frequently resort to manual segmentation due to lack of sufficient training data for deep learning methods, high cost of generating ground-truth data (Sun *et al*., 2022) and need for retraining the algorithms for different morphologies, resolution, image quality, or acquisition techniques (Meijering, 2020; Pelt, 2020). In such cases the modularized CellWalker pipeline allows the users to use such manual or hand-painted image segmentations directly for 2D/3D morphological characterization of segmented objects. The segmentation viewer module available as IPython notebook can read such manual segmentations and export a 3D reconstruction which can then be transferred directly to CellWalker’s Blender add-on (discussed further) for morphological analysis (**Fig. 1E**).

### Morphological analysis

The second module of the CellWalker pipeline is designed for morphological characterization. This open-source module runs under Blender and can be installed as a standard Blender add-on (**Fig. 1E, SI Section 2**). The graphical interface of the Blender addon is a panel located in the Sidebar which can be opened by pressing ‘N’ on keyboard.. Starting with the import of objects (.OBJ files), the addon provides functions including calculation of distance, path length, surface area, volume, and perimeter (More information is provided in the wiki pages of the repository on GitHub). Additionally, the in-build 3D-modeling functionalities of Blender remain at user’s disposal.

The Blender add-on also provides specialized functions such as skeletonization (Silversmith *et al*., 2021), cross-sectional features by slicing the objects (**Fig. 1E**) and distribution of organelles inside cells. The results of morphometric characterization are exported in a tabular format as. CSV files (**Fig. 1E**).

## Conclusion

We designed a python-based pipeline, named CellWalker, for segmentation and morphological analysis of microscopic 3D images of cells. Our pipeline bridges the segmentation protocols with 3D morphological measurements while keeping the corresponding modules independently accessible. This modular approach allows us to take advantage of existing computing infrastructures and tools such as Google Colab, IPython notebooks and the Blender software package. The design of CellWalker is also kept open-source and visible-source as much as possible to make it useful for both programmers and non-coding users.

## Author contributions

Conceptualization: HK, CZ

Code: HK, NDM

Annotation of images: HK, NDM

HK and NDM wrote the manuscript with CZ input. All authors revised and approved the final version.

## Funding

This work has been supported by the Inception program (Investissement d’Avenir grant ANR- 16-CONV-0005), the ANR-17-CONV-0005 Projet Q-life and by the Big Brain Theory Call for projects and internal seed grant from Institut Pasteur to Chiara Zurzolo.

## Acknowledgments

We thank DC Cervantes for initial manual segmentations and helpful discussions during development of the CellWalker pipeline. We also thank J. Lichtman and A. Wilson for making electron microscopy data available for testing, M. Rakotobe for providing confocal imaging data for testing, Flavia Salinas for helping with parallel fiber segmentation and Orfane Coulon-- Mahdi for identification of Golgi bodies and Centrosomes in Granule Cells for demonstration. We also thank the members of Zurzolo lab for their support and discussions.

## Notes

https://github.com/utraf-pasteur-institute/CellWalker-notebooks

https://github.com/utraf-pasteur-institute/CellWalker-blender

### Competing Interest Statement

The authors have declared no competing interest.

### Summary of Updates

Author affiliations updated.

