## Supplementary Information for "CellWalker: A user-friendly and modular computational pipeline for morphological analysis of microscopy images"

### User Guide

Instructions to install and use the CellWalker pipeline can be found online at following locations-

**CellWalker-notebooks-** https://github.com/utraf-pasteur-institute/CellWalker-notebooks/blob/main/README.md

**CellWalker-blender-** https://github.com/utraf-pasteur-institute/CellWalker-blender/wiki

### Code and data availability

CellWalker is fully open source and the code is available on GitHub in the following repositories.

https://github.com/utraf-pasteur-institute/CellWalker-notebooks

https://github.com/utraf-pasteur-institute/CellWalker-blender

### The CellWalker pipeline

The CellWalker pipeline is divided in two modules- CellWalker-notebooks and CellWalker-blender.

### Section 1: CellWalker-notebooks

There are two Jupyter notebooks included in this module.

#### Section 1.1: Automated segmentation

The notebook named **'Segmentation_CNN_UNET.ipynb'** provides a protocol for automated segmentation of microscopy images using a UNET convolutional neural network (CNN) architecture. It is recommended to run this notebook on cloud computing platforms such as Google Colab. This notebook has been tested on Google Colab.

The automated segmentation notebook is self-explanatory and the required instructions are present as markdown blocks in the notebook. The notebook walks the user through following main steps.

1. Getting started- Brief description on how to execute the notebook
2. Getting set up- Mounting Google Drive, installing and importing modules
3. Setting up Colab session- Google Colab set up
4. Input and pre-processing- Define input images and set up the images for training and testing
5. Set up data pipelines and visualization methods- Define classes and functions for data input/visualization
6. Define U-NET parameters- Set the parameters of a U-NET
7. Create model- Create a CNN U-NET model object with input parameters
8. Set up datasets and output directories- Define model validation dataset output name for trained model
9. Train the model- Fit the model to input training data
10. Plot results- Display plots for model training process
11. Model evolution- Evaluate the trained model on test images and apply on new images to get results (predicted masks)
12. Post-processing- Process predicted masks to label segmented regions.

**Demonstration: Segmentation of parallel fibers in EM images**

**Dataset used**: Serial sectioning electron microscopy data of a mouse cerebellum at P7 (Wilson *et al.*, 2019). (1.7 x 10^6 µm^3 collected at 4x4x30 nm^3 per voxel resolution)

**Raw data availability**: https://bossdb.org/project/wilson2019 (Wilson *et al.*, 2019)

**Training data (Ground truth)**: 85 slices, 400 x 400 pixel^2 tiles at 4 x 4 nm^2 per pixel resolution (mip0 resolution). Manually segmented.

Pre-processing was performed on the ground truth segmentation to shrink the segmentation masks (see notebook ?). This is an optional step which may be skipped for other examples.

**Ground truth data split for training purpose**:

Train images: 60 (70%), Validation images: 16 (20%) , Test images: 9 (10%)

**Example ground truth image and mask pairs**:

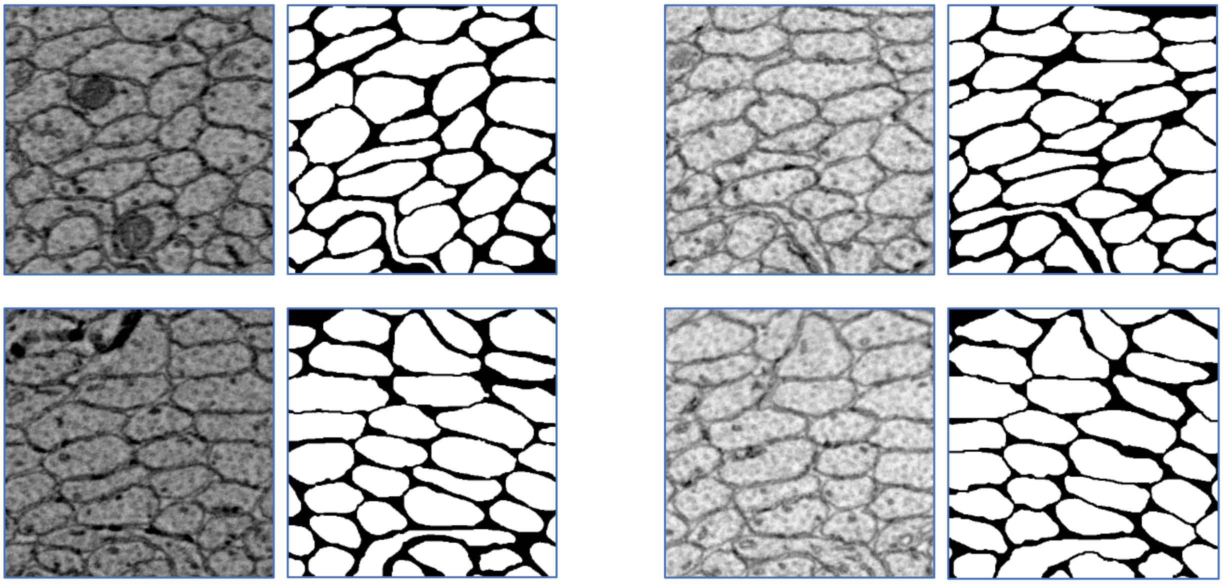

**Training performance:**

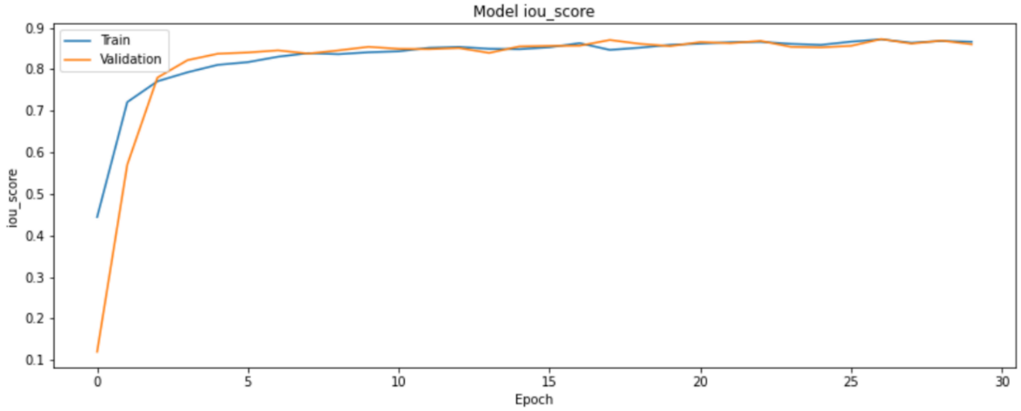


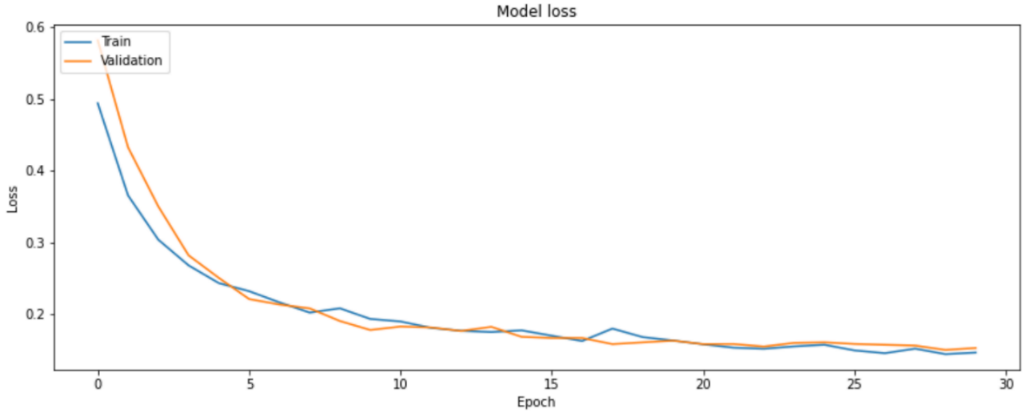

**Testing:**

Mean loss: 0.14352

mean iou_score: 0.87465

mean f1-score: 0.93307

**Inference:**

P7 mouse cerebellum data, 1001 slices, 400 x 400 pixel^2 tiles (chosen from a region different than training data)

The segments are artificially colored during post-processing step.

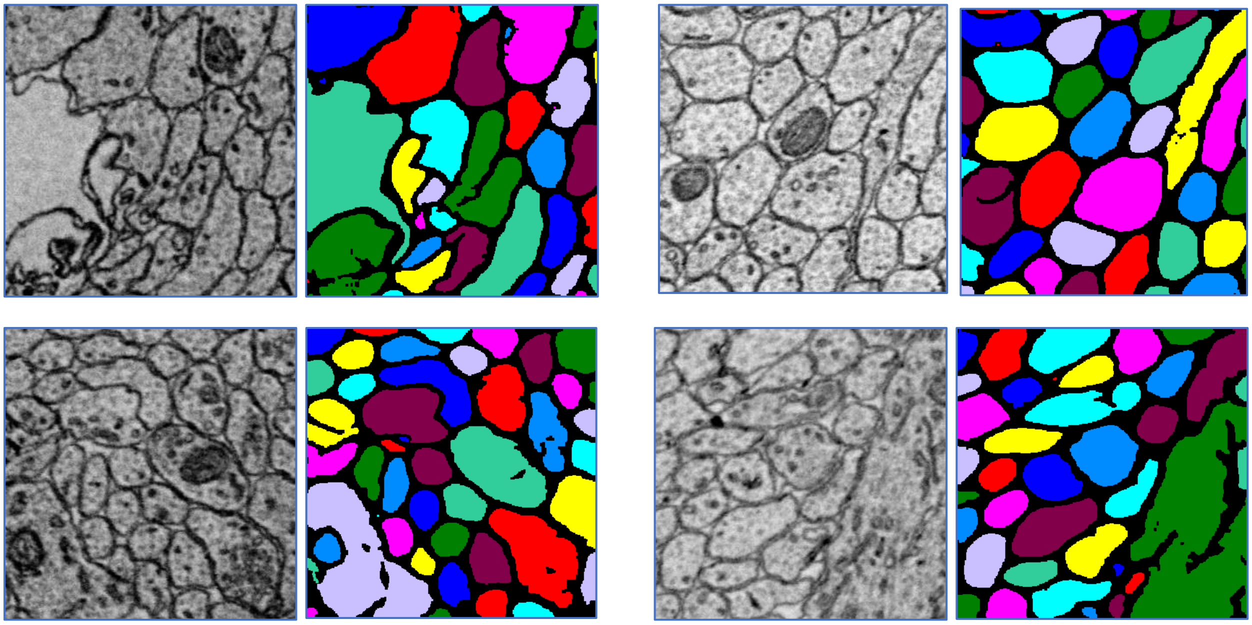

**<u>Section 1.2: Visualization of segmentation and exporting 3D objects</u>**

The notebook named '**Segmentation_visualizer.ipynb**' provides a user interface inside Jupyter-notebook environment using IPython widgets to visualize image segmentation. It allows users to interactively load raw and segmented 3D images (image stacks) side-by-side. This notebook depends on the segviewer.py and skeletontools.py files provided in the repository. While the main functionalities of the code are in these two .py files, the IPython notebook provides control in a graphical interface.

Once loaded, the segments identified in the image stack are displayed on the right hand side while the scrollable image stacks are shown on the left hand side panel. For multiple segments users can also assign colors to the segments for visual analysis.

The selected segments can be exported to a .OBJ file which serves as an input for the next module in the CellWalker pipeline, CellWalker-blender, for 3D morphometric analysis.

**Demonstration:**

Visualization of cropped segmented EM data from P7 mouse cerebellum- A tunneling connection between granule cells

**Segmentation visualizer can open raw and segmented image stacks side-by-side**

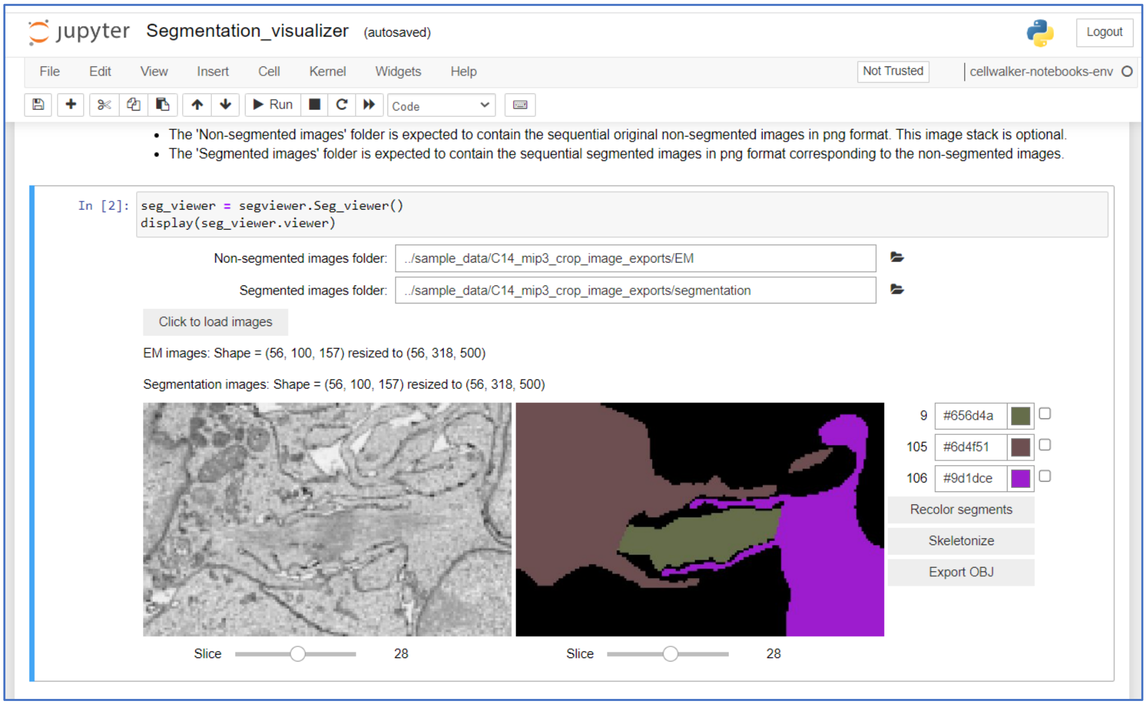

### Segment colors may be changed for better visualization

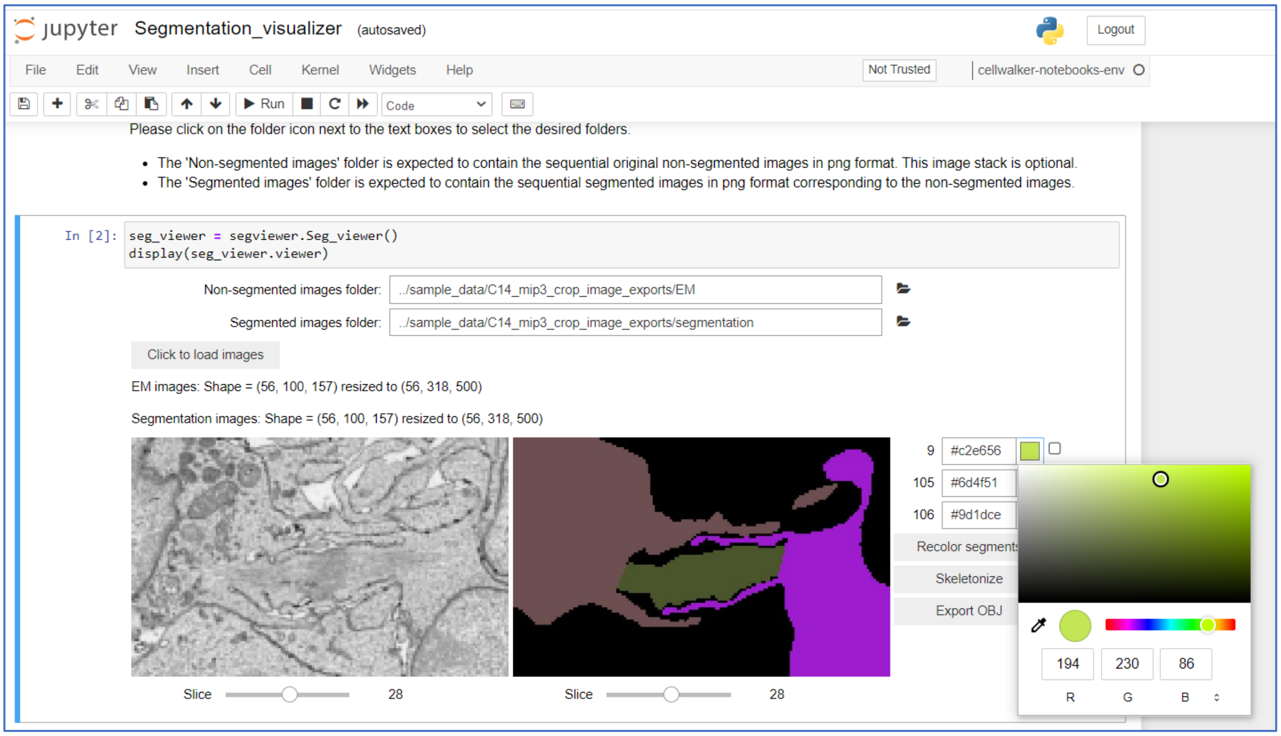


### Color of segment 9 updated

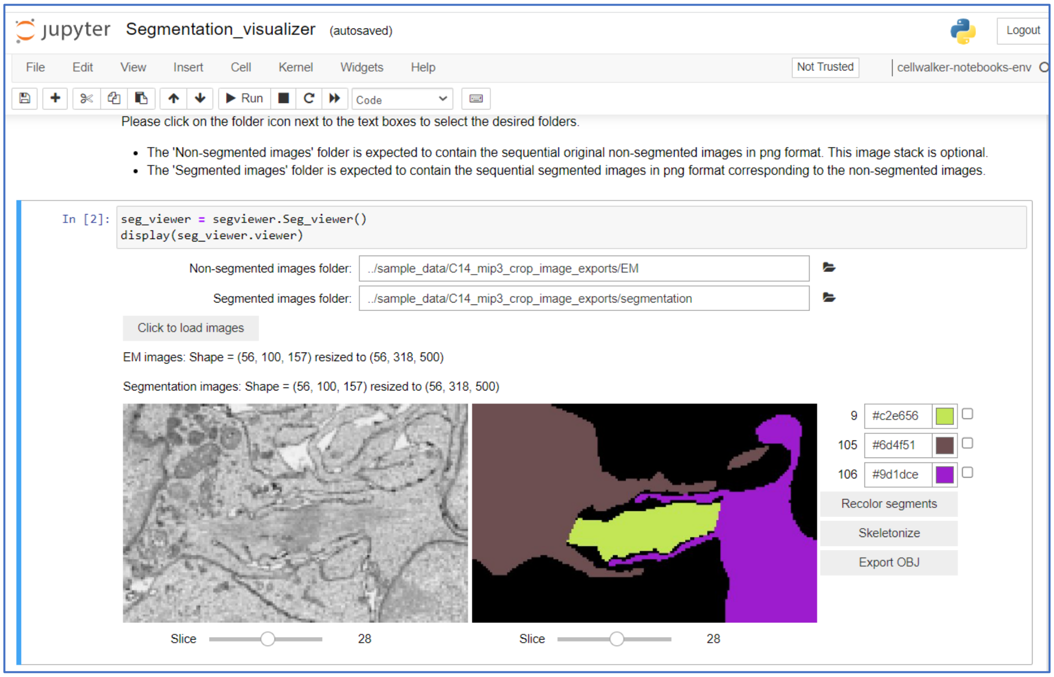

### Selected segments can be exported as .OBJ files

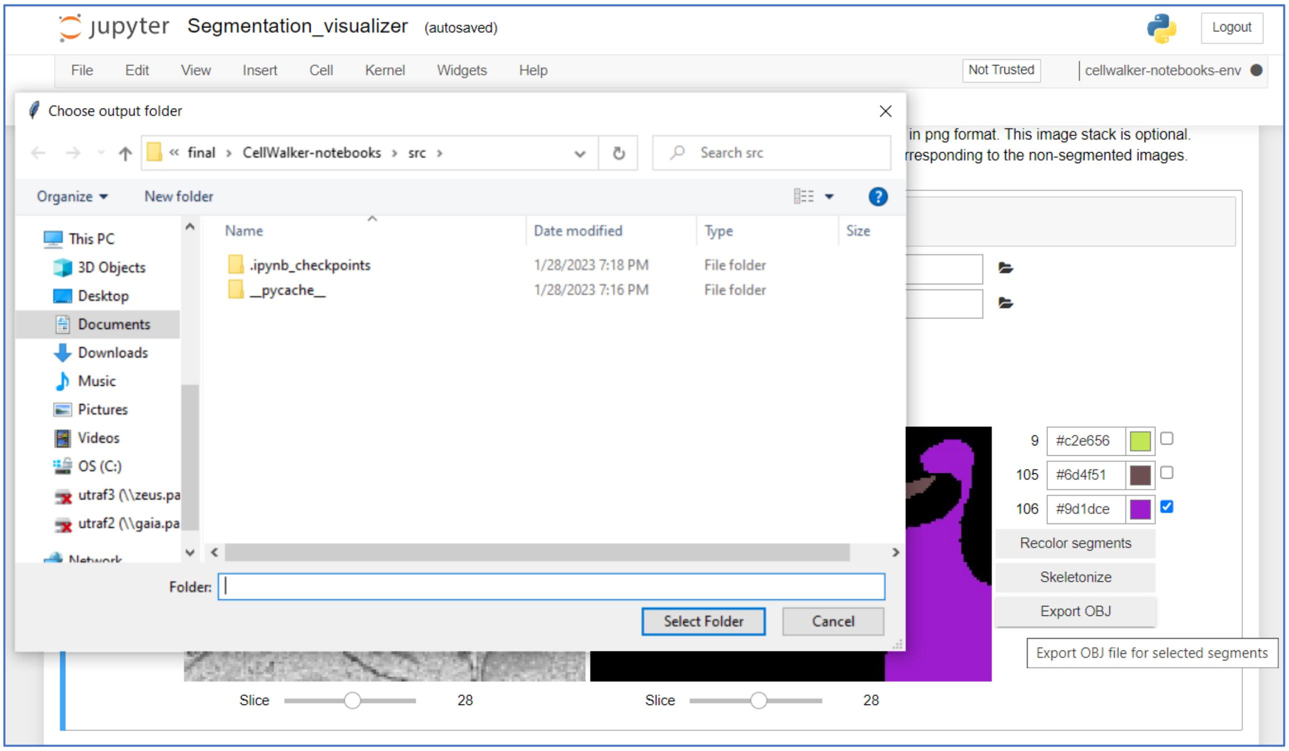


### Exported .OBJ file can be imported in 3D graphics softwares such as Blender

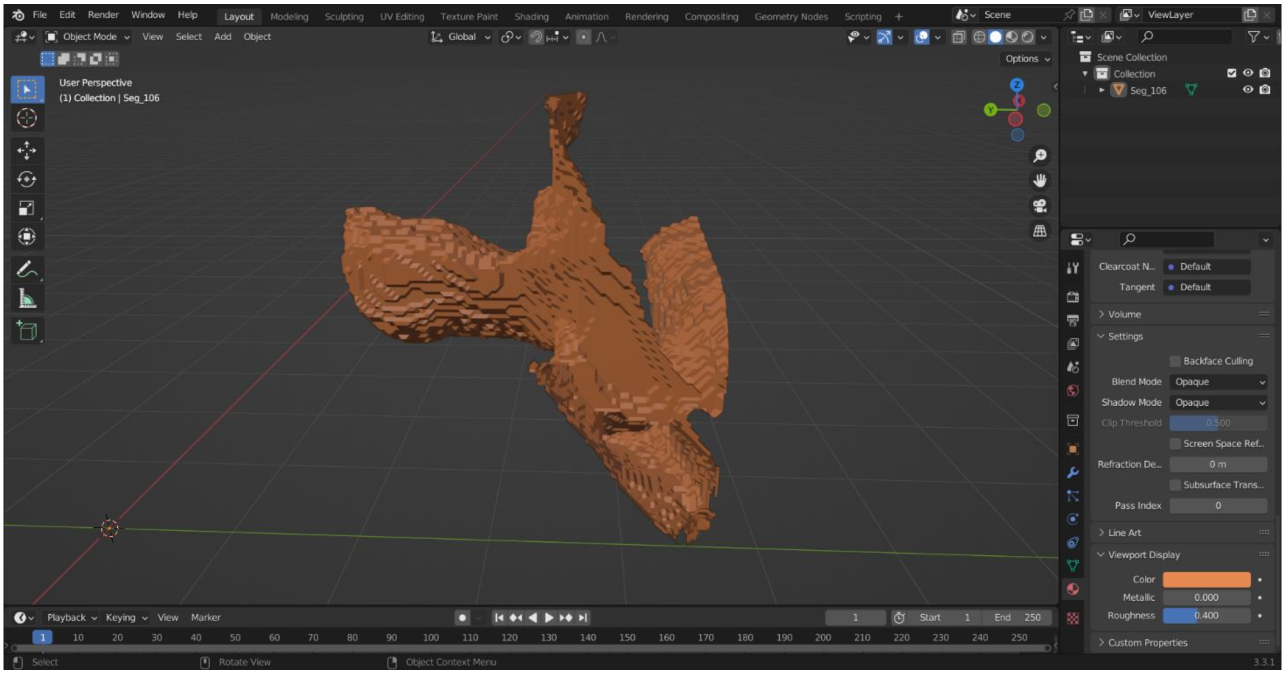

### Section2: CellWalker-blender

This is the second part of the CellWalker pipeline. CellWalker-blender is an add-on that works inside the open-source 3D modeling software, Blender. It allows the user to load the .OBJ files exported by the previous module in the CellWalker pipeline- Segmentation visualizer. The CellWalker's Blender add-on can also work on 3D objects from other sources, however it has not been tested on such externally sourced 3D objects.

Below are the functionalities provided by CellWalker-blender for morphometric analysis of 3D models of biological cells and organelles. More detailed descriptions and installation/usage instructions can be found on GitHub (https://github.com/utraf-pasteur-institute/CellWalker-blender/wiki)

1. Import .OBJ files: Imports multiple .OBJ files in a folder at once. Note that it is also possible to import the .OBJ files one at a time using Blender's File > Import menu.
2. Surface area and volume: Calculates surface area and volume of multiple selected objects and saves to a .CSV file in selected folder.
3. Cross-section tool: Creates cross-sections at specified distance between consecutive slices along a chosen direction, also calculates morphological properties of the corss-sections .
4. Distance tool: Calculates straight distance as well as mesh distance (using Dijkstra's algorithm) between selected vertices on an object.
5. Skeletonize: Builds a skeleton of a selected object using Kimimaro algorithm (Silversmith *et al.*, 2021) and allows to save the skeleton as .OBJ file.
6. Angular distribution: Calculates distribution of an organelle around the centroid of a cell, where the organelle and the cell are represented by two separate .OBJ files.

**Demonstration:** Potentially dividing granule cells in ssSEM 3D data of mouse cerebellum at P7

### Granule Cell pair and organelles loaded in Blender, with CellWalker add-on opened on right hand side of the scene

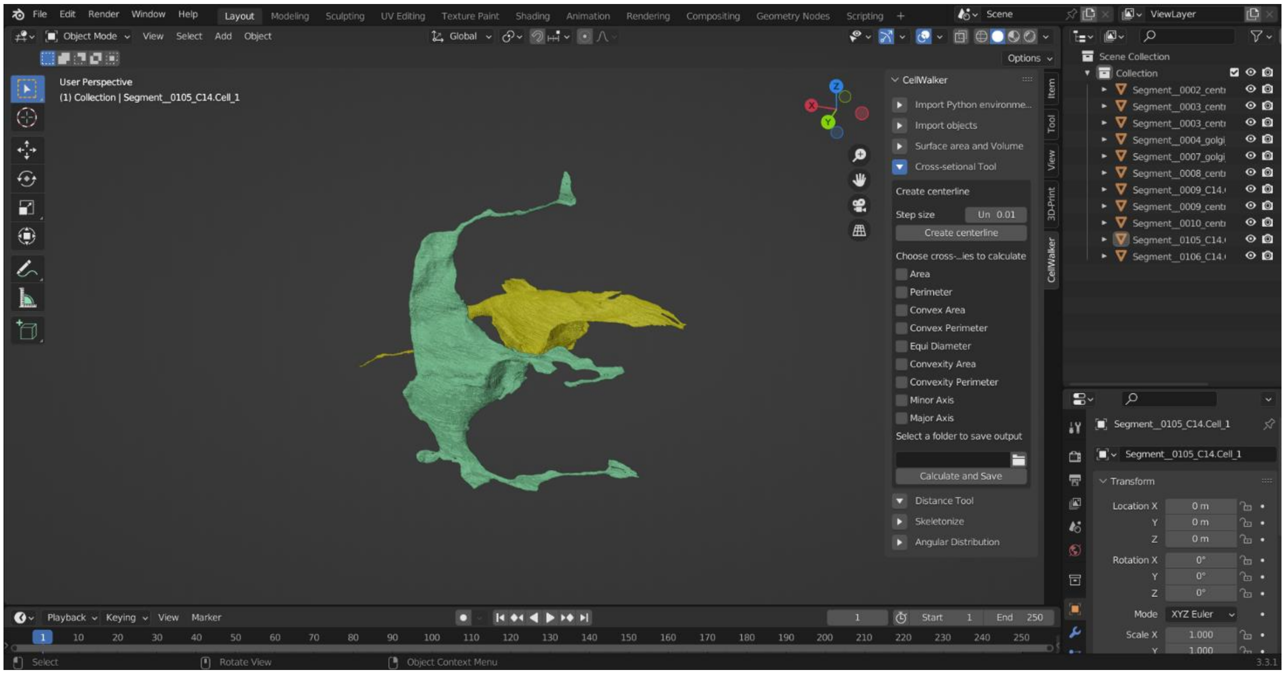


### Surface area and volume

Granule cells:

1. Cell 1: Surface area 459.5 um^2, Volume 178 um^3
2. Cell 2: Surface area 539.4 um^2, Volume 192.5 um^3

**Cross-sections and properties of cross-sections**

Cross-sectioning performed on a lamellipodium of Cell 1.

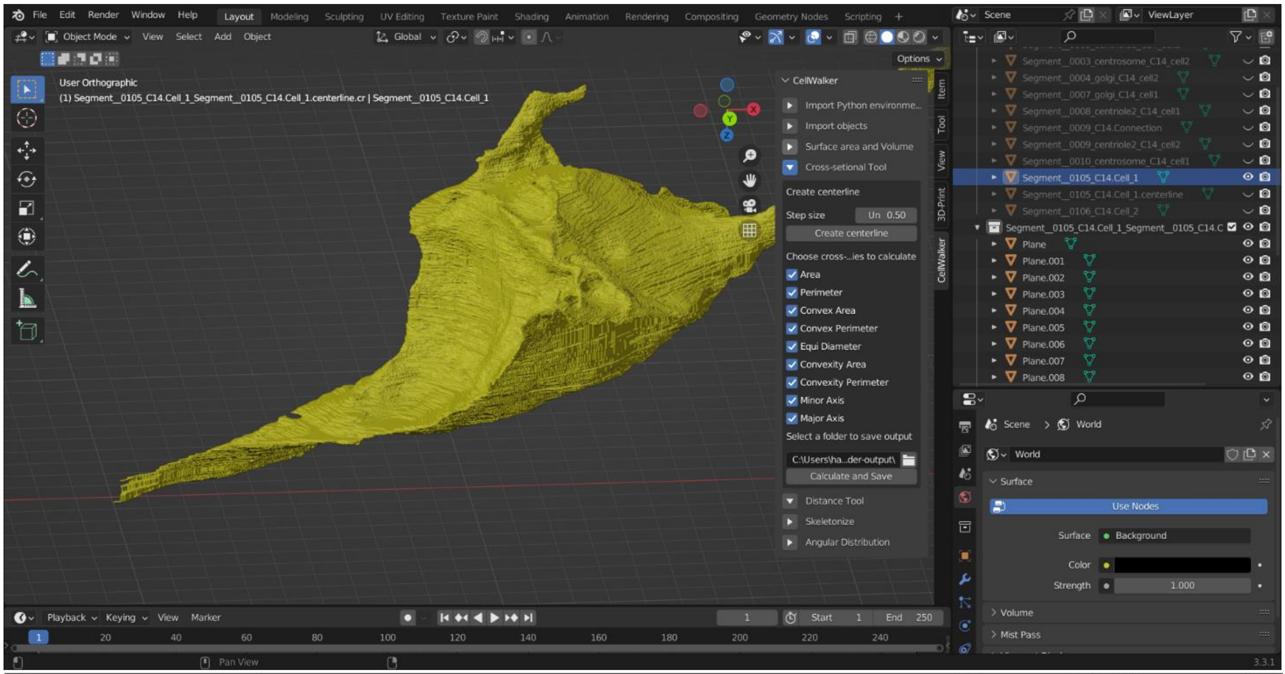


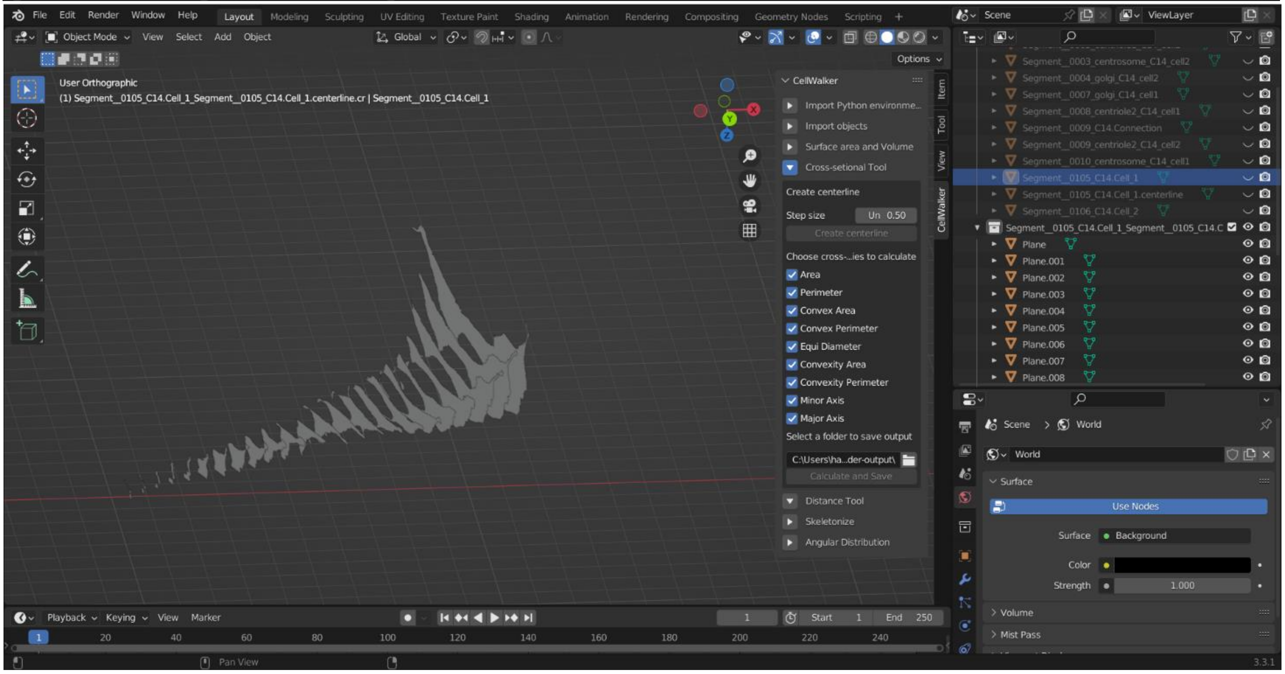

**Properties of cross-sections exported as a CSV file**

| IND | Area | Perimeter | Convex Area | Convex Perimeter | Equiv Diameter | Convexity _area | Convexity _perimeter | Minor Axis | Major Axis |
|---|---|---|---|---|---|---|---|---|---|
| **0** | 12.31 | 30.45 | 20.62 | 20.62 | 3.96 | 0.60 | 0.68 | 4.71 | 7.76 |
| **1** | 8.61 | 24.42 | 17.80 | 17.80 | 3.31 | 0.48 | 0.73 | 3.63 | 6.90 |
| **2** | 5.65 | 23.07 | 16.62 | 16.62 | 2.68 | 0.34 | 0.72 | 3.14 | 6.73 |
| **3** | 3.97 | 46.58 | 12.02 | 12.02 | 2.25 | 0.33 | 0.26 | 2.00 | 5.01 |
| **4** | 2.79 | 15.97 | 10.63 | 10.63 | 1.88 | 0.26 | 0.67 | 1.79 | 4.48 |
| **5** | 2.28 | 538.68 | 10.90 | 10.90 | 2.82 | 0.57 | 0.02 | 1.65 | 4.33 |
| **6** | 1.77 | 13.36 | 9.34 | 9.34 | 1.50 | 0.19 | 0.70 | 1.41 | 4.19 |
| **7** | 1.62 | 13.30 | 8.82 | 8.82 | 1.44 | 0.18 | 0.66 | 1.24 | 3.94 |
| **8** | 1.69 | 13.35 | 8.65 | 8.65 | 1.47 | 0.20 | 0.65 | 1.27 | 3.70 |
| **9** | 1.41 | 9.73 | 8.23 | 8.23 | 1.34 | 0.17 | 0.85 | 1.18 | 3.64 |
| **10** | 1.12 | 8.87 | 7.98 | 7.98 | 1.19 | 0.14 | 0.90 | 1.17 | 3.53 |
| **11** | 1.03 | 7.18 | 6.34 | 6.34 | 1.14 | 0.16 | 0.88 | 1.02 | 2.81 |
| **12** | 1.00 | 7.31 | 6.25 | 6.25 | 1.13 | 0.16 | 0.86 | 1.06 | 2.76 |
| **13** | 1.01 | 6.78 | 5.88 | 5.88 | 1.13 | 0.17 | 0.87 | 0.96 | 2.65 |
| **14** | 0.97 | 6.56 | 5.49 | 5.49 | 1.11 | 0.18 | 0.84 | 0.91 | 2.46 |
| **15** | 0.91 | 6.08 | 5.36 | 5.36 | 1.08 | 0.17 | 0.88 | 0.91 | 2.40 |
| **16** | 0.73 | 5.15 | 4.53 | 4.53 | 0.97 | 0.16 | 0.88 | 0.83 | 1.95 |
| **17** | 0.48 | 4.30 | 3.76 | 3.76 | 0.78 | 0.13 | 0.87 | 0.65 | 1.63 |
| **18** | 0.21 | 3.54 | 3.09 | 3.09 | 0.51 | 0.07 | 0.87 | 0.35 | 1.43 |
| **19** | 0.14 | 2.92 | 2.67 | 2.67 | 0.43 | 0.05 | 0.91 | 0.33 | 1.22 |
| **20** | 0.09 | 2.27 | 2.07 | 2.07 | 0.33 | 0.04 | 0.91 | 0.30 | 0.92 |
| **21** | 0.05 | 1.94 | 1.74 | 1.74 | 0.26 | 0.03 | 0.90 | 0.39 | 0.70 |
| **22** | 0.04 | 1.73 | 1.51 | 1.51 | 0.24 | 0.03 | 0.87 | 0.32 | 0.61 |

Trends in Area and Major axis of along the cross-sections indicates tapering nature of the lamellipodium.

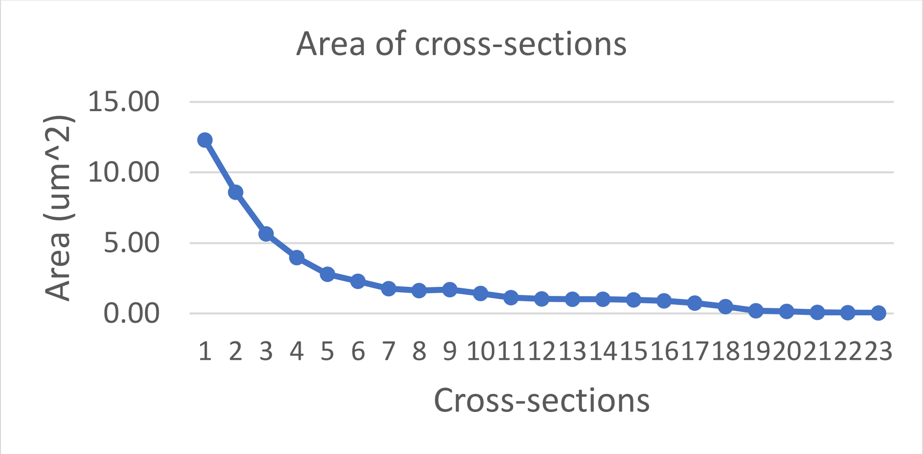


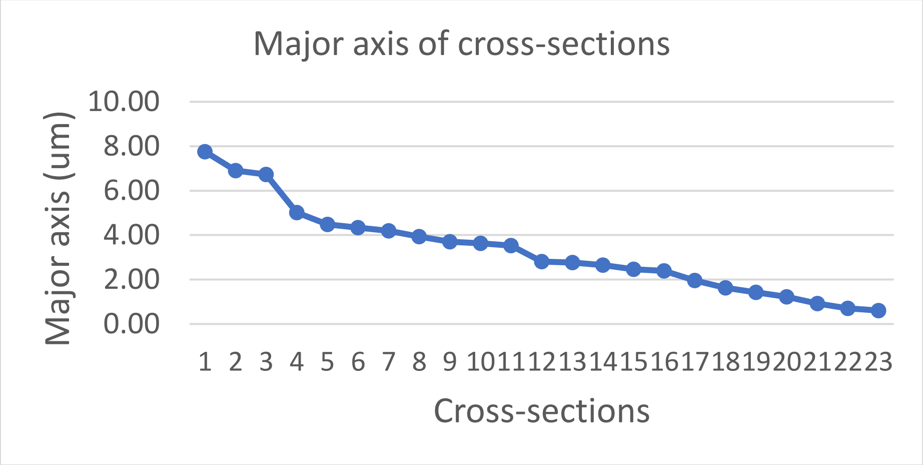


**Distance calculation**

Straight vs Dijkstra distance (topological distance) between ends of two lamellipodia of Cell 2
Note: Calculation of Dijkstra distance over large cells can take long time ( few minutes). It is not advisable to calculate topological distances on cells over long distances unless actually useful. A faster alternative is to compute Dijkstra distance on a skeleton of a cell.

The following figures display the process of skeletonization and distance calculation.

Cell 1

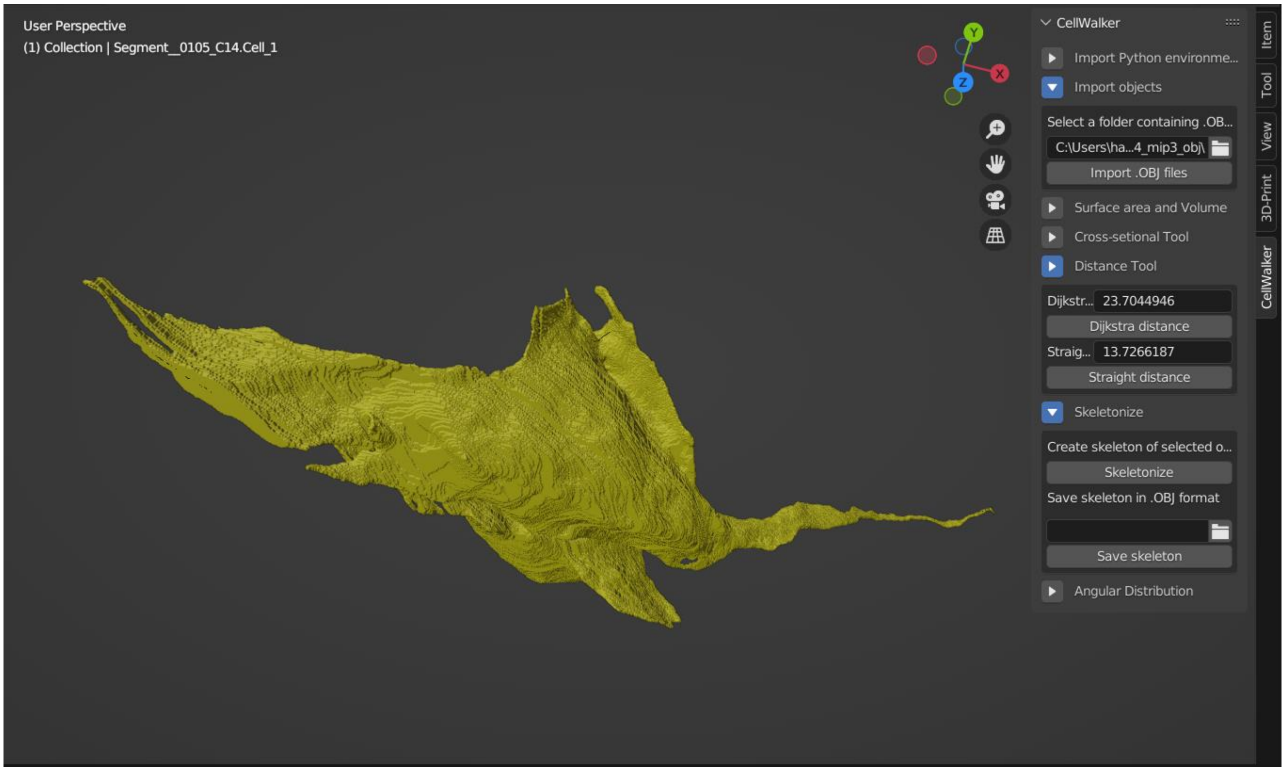


Skeletonized Cell 1

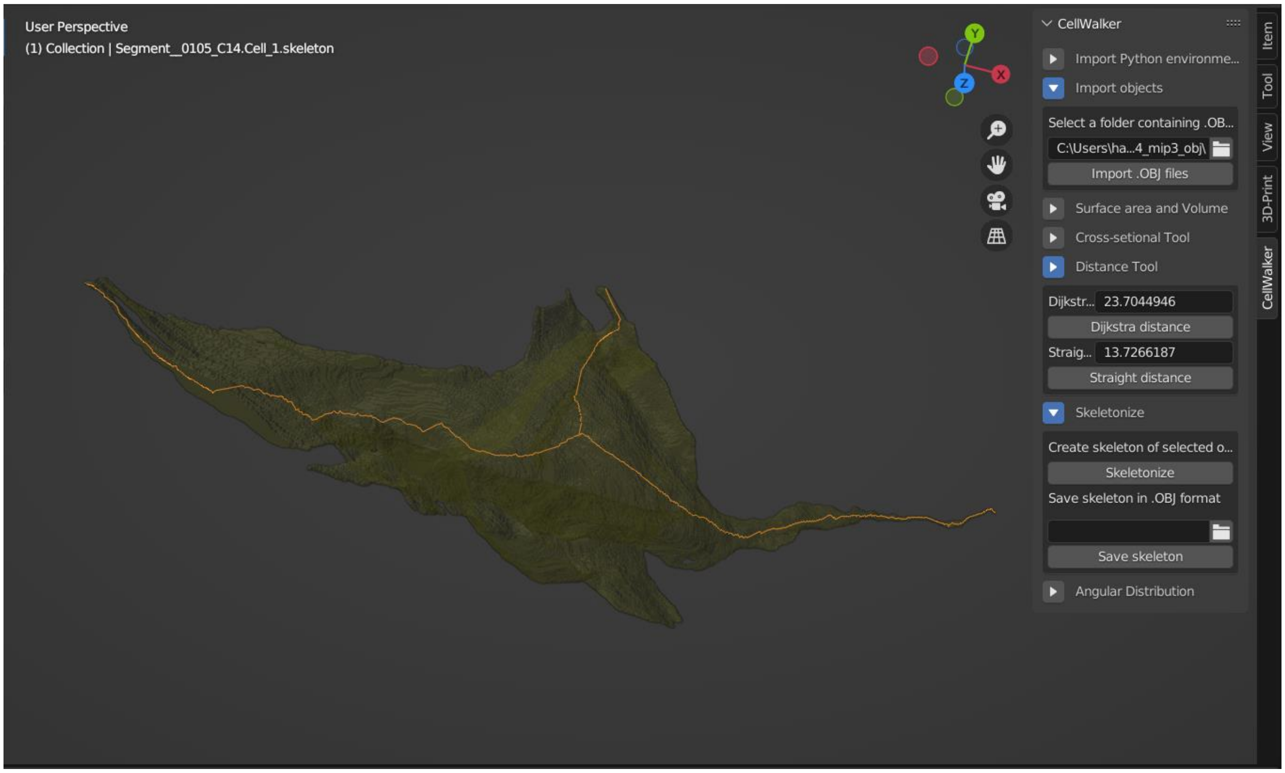

Straight distance between two lamellipodia using skeleton (13.7 um)

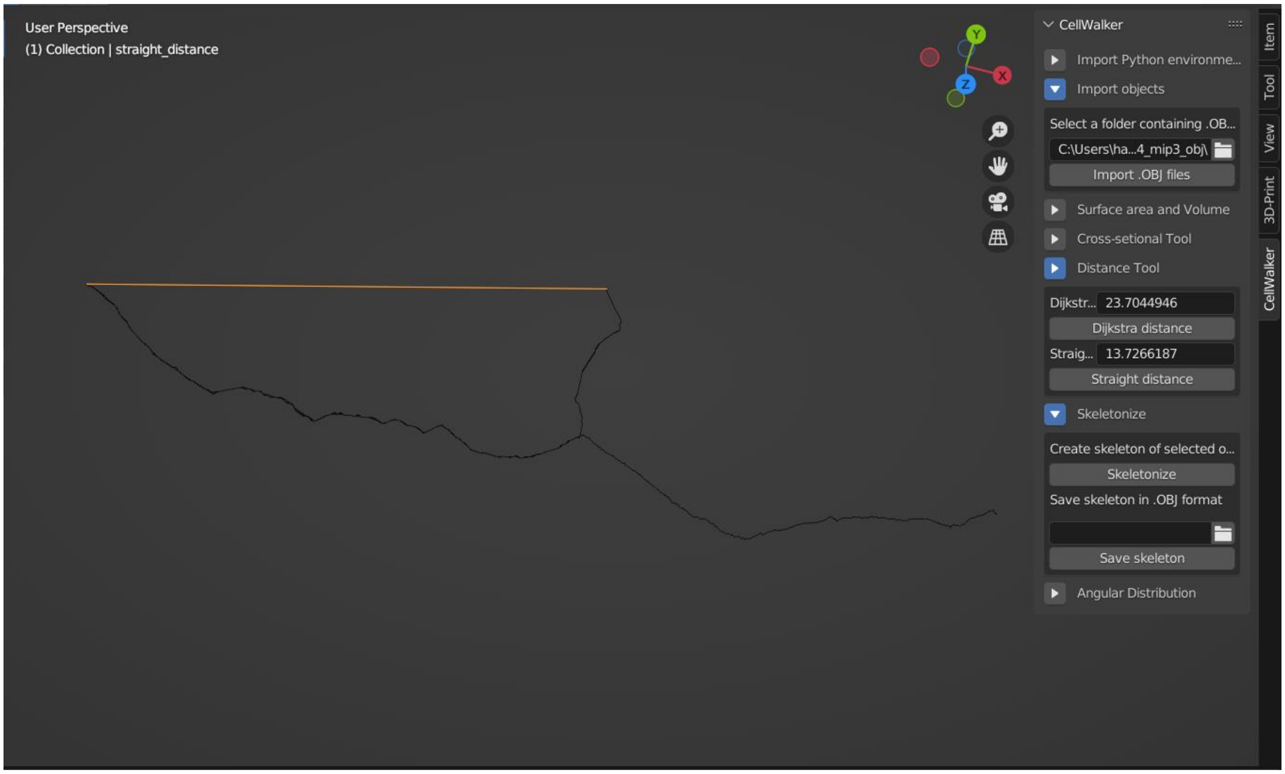


Dijkstra distance (topological distance) between two lamellipodia using skeleton (23.7 um)

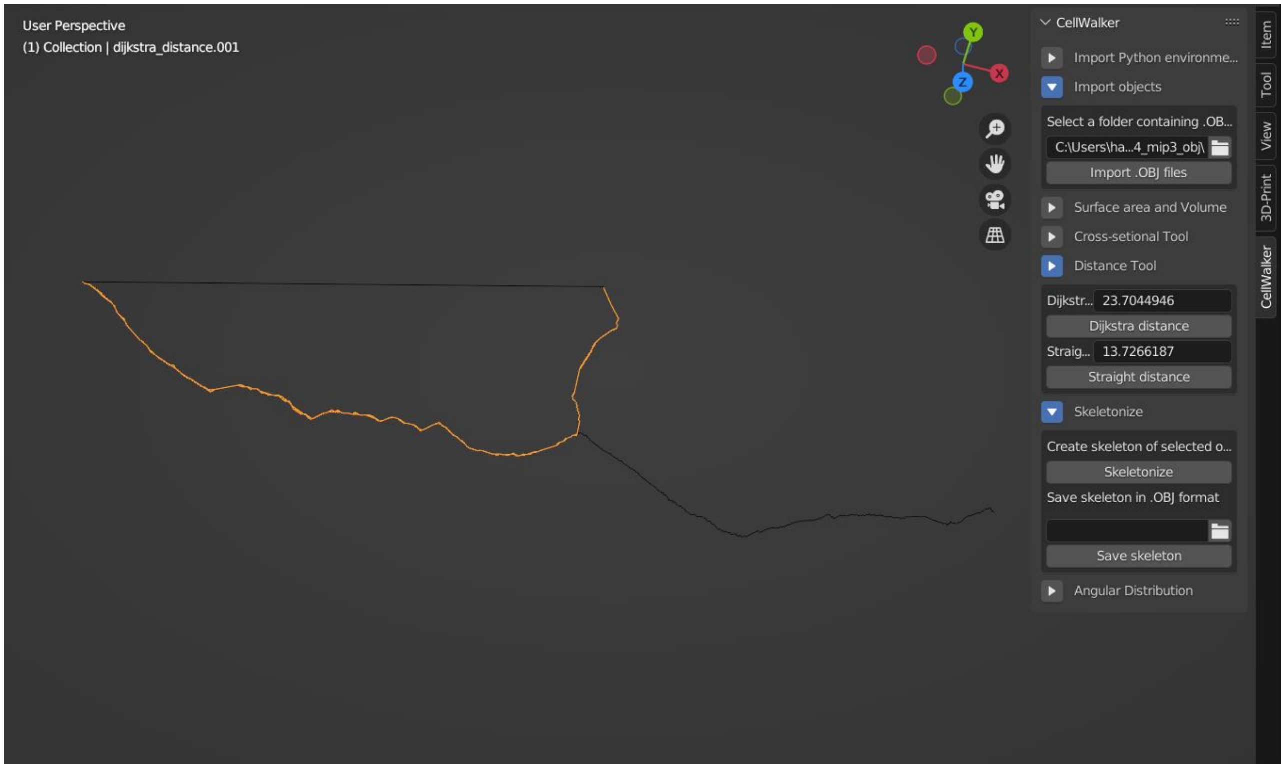

**Distribution of Golgi bodies in Cell 1**

Mean angular distribution: 65.8 degrees

Standard deviation in angular distribution: 34.14

Mean angular distribution less than 90 degrees indicates that the Golgi bodies could be spread across the cell to a smaller extent.

(Very high values of mean angular distribution (close to 180 degrees) would have indicated a uniform spread of Golgi bodies across the cell.)

Golgi bodies shown in Cyan color in the following figure

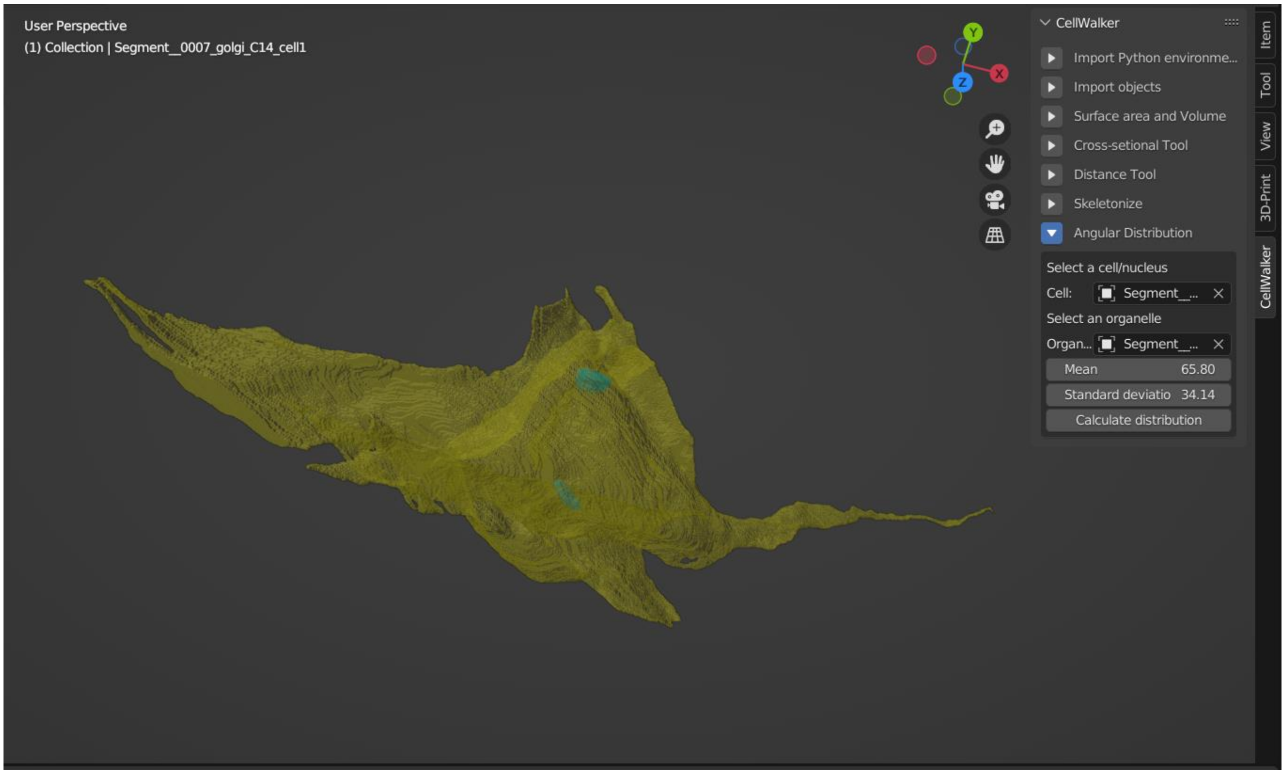
